# Breast cancer dormancy is associated with a 4NG1 state and not senescence

**DOI:** 10.1101/2020.11.20.367698

**Authors:** Chloé Prunier, Ania Alay, Michiel van Dijk, Kelly L. Ammerlaan, Sharon van Gelderen, Dieuwke L. Marvin, Amina Teunisse, Roderick C. Slieker, Karoly Szuhai, A.G. Jochemsen, Xavier Solé, Peter ten Dijke, Laila Ritsma

**Affiliations:** Department of Cell and Chemical Biology and Oncode Institute, Leiden University Medical Center, 2333 ZC Leiden, the Netherlands; Oncode Institute, the Netherlands; R&D department, Inovotion SAS, 38700 La Tronche, France; Unit of Bioinformatics for Precision Oncology, Catalan Institute of Oncology, L’Hospitalet de Llobregat, Spain; Molecular Mechanisms and Experimental Therapy in Oncology Program (Oncobell), Bellvitge Biomedical Research Institute (IDIBELL), L’Hospitalet de Llobregat, Spain; CIBER of Epidemiology and Public Health (CIBERESP), Madrid, Spain

**Keywords:** breast cancer, cell cycle arrest, cellular dormancy, fucci, liver metastasis, senescence

## Abstract

Reactivation of dormant cancer cells can lead to cancer relapse, metastasis and patient death. Dormancy is a non-proliferative state and is linked to late relapse and death. No targeted therapy is currently available to eliminate dormant cells, highlighting the need for a deeper understanding and reliable models. Here, we thoroughly characterize the dormant D2.OR and proliferative D2A1 breast cancer cell line models *in vivo* and *in vitro*, and assess if there is overlap between a dormant and a senescent phenotype. We show that D2.OR but not D2A1 cells become dormant in the liver of an immunocompetent model. *In vitro*, we show that D2.OR cells are polyploid ER^+^/Her2^+^ cells, and in response to a 3D environment are growth arrested in G1, of which a subpopulation resides in a 4NG1 state. The dormancy state is reversible, and not associated with a senescence phenotype. This will aid future research on breast cancer dormancy.

## INTRODUCTION

Breast cancer (BC) is the second most-deadliest cancer in women worldwide^1^. Most BC patients do not succumb to the primary tumor, but rather die from overt metastases that arise later during disease progression. Follow up analysis showed that 62% of BC patients die 5 to 20 years after initial treatment due to late relapse (recurrence or metastasis), particularly patients with luminal (estrogen receptor (ER^+^); human epidermal growth factor receptor (Her2^+^)) breast cancer. Late relapse has been linked to a non-proliferative cell state called dormancy^2^. Dormant cells can remain undetected in the tissue for years or decades, until their escape results in proliferation and the formation of recurrent disease or metastasis. In addition, dormancy is at the basis of a drug tolerance state in certain models, allowing cells to evade apoptosis and survive anti-cancer therapy^3–5^. As such, dormancy is a major clinical issue as reactivation of dormant cancer cells can lead to cancer relapse and eventually patient death^6,7^.

Several signaling pathways have been described to promote dormancy such as the transforming growth factor beta (TGF-β) family^8^, the urokinase receptor uPAR and downstream targets p38α/β and ERK1/2 MAP kinases, the cyclin-dependent kinase inhibitors (CDKN) 1A (p21) and 1B (p27), integrins^9^, and others reviewed here^8,10,11^. Most of these pathways regulate proteins involved in the cell cycle, and ultimately result in a G0/G1 cell cycle arrest. As a consequence, dormant cells are resistant to chemotherapies that target proliferating cells^4,5^. Moreover, no targeted therapy is currently available in the clinic to eliminate dormant cells, nor are reliable prognostic markers that can predict late relapse. This highlights the need for an even deeper understanding of the signaling pathways that govern the dormancy process.

Senescence, a physiological response to replicative or oncogenic stress in normal cells, shares characteristics with dormancy. Similar to dormancy, senescent cells exit the cell cycle and do not proliferate^12^. Senescence was considered irreversible for decades, but in the last 10 years reversible senescence named “pseudo-senescence” or “senescence-like” has been observed and documented^13,14^. The mechanisms underlying senescence are quite well defined and a variety of markers have been identified. Senescent cells are often characterized by DNA damage, expression of senescence activated beta-galactosidase (SA-βGal), loss of Lamin B1, and secretion of a senescence-associated secretory profile (SASP). The program can be induced upon activation of tumor suppressor p53 or upregulation of cyclin-dependent kinase inhibitor CDKN2A (p16)^12^. Interestingly, some dormant cells were shown to have senescence phenotypes like SASP and SA-βGal ^3,15^.

To gain insight into signaling pathways that regulate dormancy and to better understand a possible role for a senescence program in dormancy, proper dormancy models are required. For breast cancer, only a few models exist, which can be used both *in vivo* and *in vitro* (e.g. MCF7). The most commonly used model being the D2.OR murine mammary cancer cell line, which is then frequently compared side-by-side to the related D2A1 (D2) proliferative tumor cell line^9^. Both cell lines were derived from different D2 hyperplastic alveolar nodule mammary tumors^16–18^. *In vivo*, D2.OR cells disseminate, but remain dormant, whereas D2A1 cells form metastases^17,19,20^. This can be modeled by the Matrigel on Top assay (MoT), in which cells are placed on a thin 3D layer of growth factor-reduced Matrigel. In this model, the D2.OR cells remain dormant, whereas the D2A1 cells do not remain dormant but proliferate^21^. The D2A1 cells were initially characterized as triple negative, but other studies have suggested that they were ER^+^ or luminal B^22,23^. The molecular subtype of the D2.OR cells has not been assessed in the literature. As the field of dormancy is expanding^24^ and this model is increasingly used in literature^9,25–28^, we felt the need to thoroughly characterize this D2.OR model, and assess if there is overlap with a senescence phenotype.

## RESULTS

### D2 murine mammary cancer cells are ER^+^/Her2^+^ Basal-like cells

D2.OR and D2A1 cells are often used to study tumor progression, as they are related tumor cell lines which differ in their ability to form metastases^29^. As the cells are derived from the same tumor type, but were derived from separately formed tumors in different mice, we wondered how related the cell lines were. We used RNA-seq to assess the differences and commonalities between these cells at the transcriptomic level and identified 1619 differentially expressed genes, of which 522 were downregulated and 1097 were upregulated (**Fig 1a**). We performed gene set enrichment analyses (GSEA) on the gene expression differences between the D2.OR and the D2A1 cell line in 2D culture conditions. Significantly differentially enriched gene sets (FDR<0.05) were grouped according to their degree of gene overlap (**Supplementary Fig 1a**). All significantly enriched gene set categories (e.g. related to phospholipase C (PLC) activity, cell migration and adhesion, inflammatory response and embryonic development) were upregulated in the D2.OR cells (i.e., positive Normalized Enrichment Score). Combined, these results suggest that the D2.OR cell line when compared to D2A1 cell line shows upregulation of specific pathways and cellular functions that are mostly related to cell signaling and development, as well as inflammatory response processes. A complete list of gene sets tested and their results is available in Supplementary table 1.

**Fig. 1:**
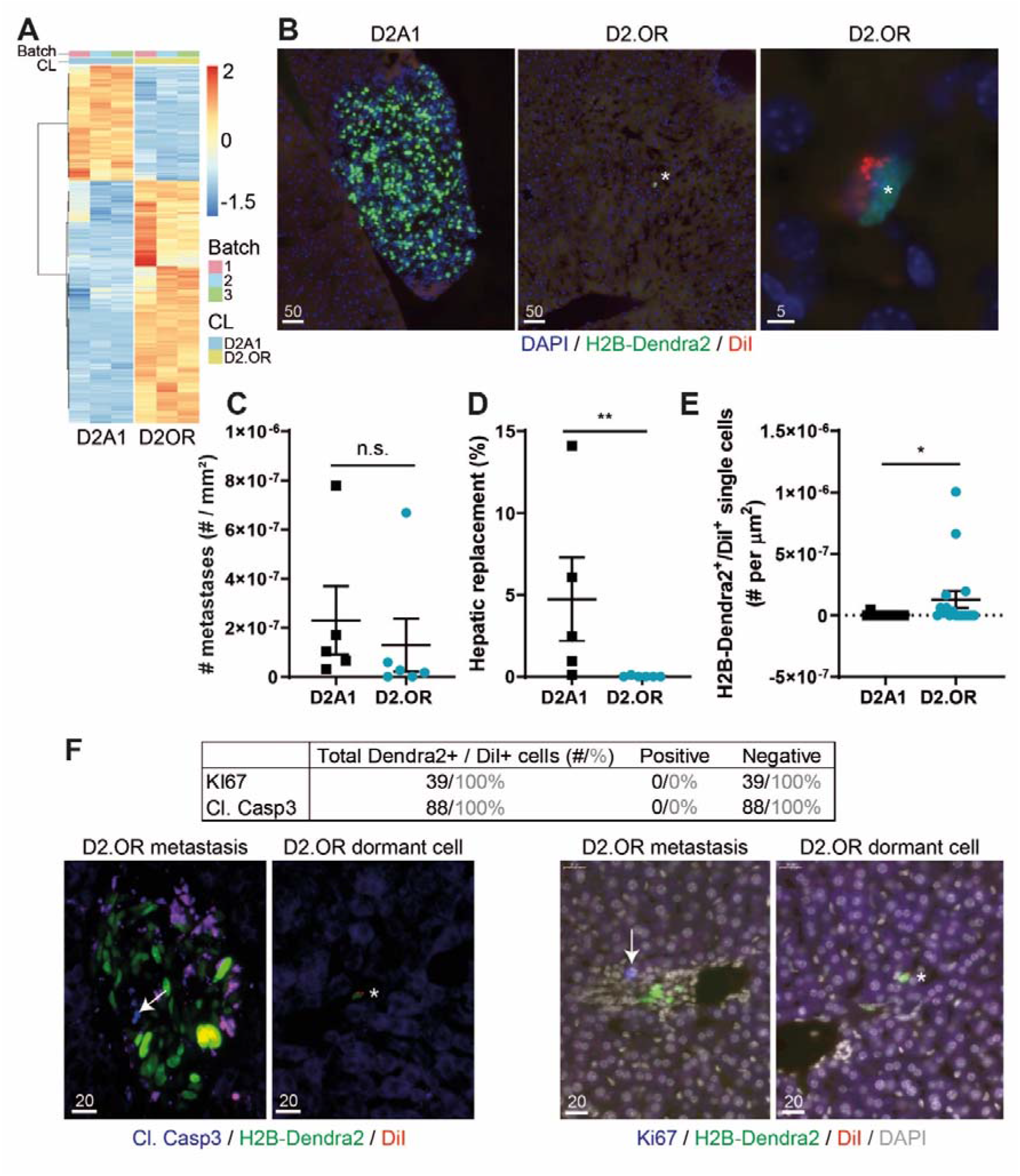
Dormancy of the D2.OR cells in liver parenchyma of syngeneic mice. **a** Heatmap of differentially expressed genes between D2.OR and D2A1 cells cultured in 2D. A total of 1619 genes were selected, based on FC > 1.5, no batch inconsistencies, and a Bonferroni correction for multiple testing > 0.05. CL = cell line. Batch = replicate. **b** Microscopy images of D2A1 or D2.OR cell (clusters) in the liver of mice 2 weeks after intramesenteric vein (i.mes) injection. Scalebar, μm. Analysis of **c** metastases number (including single cells), **d** hepatic replacement by tumor cells, or **e** dormant cells in liver sections of mice sacrificed 2 weeks after i.mes injection. n ≥ 5 mice, 3 sections per mouse. **f** Quantification (black: number of cells, grey: percentage) of immunostaining on dormant cells in sections with D2.OR cells. Below, representative images of D2.OR metastases and dormant cells (indicated by asterisk) stained for cleaved caspase 3 (left) or KI67 (right). Staining is indicated in blue (arrow). Scalebar, μm. N = 6 mice, 3 sections per mouse. Indicated P values were calculated using Mann-Whitney tests.

As BC molecular subtype is an important determinant for dormancy^6^ and no consensus on the status of D2.OR or A1 cells has been reached in literature^22,30^, we used our transcriptomic data to assign an intrinsic subtype. *Esr1*, encoding *estrogen receptor α*, was only expressed in D2.OR, but at very low levels (Log2TPM ~2.0). In contrast, *ErbB2* (Her2) was expressed in both cell lines, although higher in D2OR (**Supplementary Fig 1b**). In order to obtain further insight, we decided to use UMAP (user modeling, adaptation and personalization), a non-linear dimension reduction technique that projects multiple observations in a 2-dimensional space preserving the essential topological structure of the original data. A combined UMAP analysis of the D2A1 and D2.OR cell lines with TCGA primary breast tumors with available PAM50 information and using a mouse-derived intrinsic subtype 1841-gene signature^31^ suggested that the cells may belong to the basal subtype (**Supplementary Fig 1c**). Basal-like tumors consist for only 2.7% of ER+/HER+ tumors, and for 18.2% of

ER+/HER2-tumors^32^. As the expression of *ER* was quite low, we assessed if the D2 cells have functional ER protein. Upon activation by β-estradiol, ERα stimulates cell growth, and its antagonist 4-hydroxytamoxifen (4OHT) reduces cell growth. Indeed, we observed reduced proliferation when cells were stimulated with 4OHT (**Supplementary Fig 1d**), confirming the presence of ER in the cells. We conclude that our cells are assigned to the Basal-like molecular subgroup of breast cancers, but that they are positive for ER and ErbB2.

### D2.OR murine mammary cancer cells are growth arrested in the liver

The metastatic (or dormant) capacity of the D2 cells has been assessed using spontaneous and experimental liver and lung metastasis models^19,25,33^. However, liver metastases were not characterized in an immune competent model. Thus, we assessed if the D2.OR cells underwent dormancy in fully immune competent BALB/c mice using an experimental liver metastasis assay. Fluorescently labelled D2 cells were injected in the mesenteric vein and the number and amount of liver metastases (based on fluorescent H2B-Dendra2 signal) was assessed by scoring of fixed-frozen liver sections collected 14 days after implantation (**Fig 1b**). Although the same amount of lesions were identified in D2.OR and D2A1 injected mice (**Fig 1c**), the size of the D2.OR lesions was significantly lower compared to the D2A1 lesions, suggesting a growth arrest of D2.OR *in vivo* (**Fig 1d**). We performed label retention using a fluorescent dye to assess dormancy^34^. The injected cells were labeled with a red dye (DiI), which dilutes more or less equally among the two daughter cells upon cell division (**Supplementary Fig 2a**). When cell division is reduced by lowering serum levels, DiI is retained (**Supplementary Fig 2b**). Thus, single H2B-Dendra2^+^ cells that retained the dye (DiI^+^) did not divide and were considered dormant. Importantly, DiI does not affect cell proliferation *in vitro* (**Supplementary Fig 2c**). We observed very few H2B-Dendra2^+^/ DiI^+^ cells in the D2A1 model, and significantly more in the D2.OR model (**Fig 1e**). Dormancy of these single D2.OR cells was further validated by absence of proliferation marker Ki67 and Cleaved-Caspase3 (CC3) immunostaining, indicating that single H2B-Dendra2^+^/ DiI^+^ D2.OR cells are in cell cycle arrest and not apoptotic (**Fig 1f**). We then studied the *in vivo* behavior by intravital microscopy. Two weeks after injection, we identified dividing and apoptotic D2A1 cells in large metastases, and label-retaining dormant D2.OR cells (**Supplementary Fig 3**). In three hour movies we detected limited D2A1 cell migration, and no D2.OR cell migration (**Supplementary Fig 3**).

Taken together, the D2A1 cells proliferate and form metastases in the liver of fully competent BALB/c mice, whereas the D2.OR cells do not proliferate and remain in the liver mostly as single dormant cells.

### Dormant D2.OR are in G0/G1 cell cycle arrest

Dormancy is a complex and dynamic process which is more easily studied *in vitro*. Barkan *et al* showed that when the D2 cells were cultured in a 3 dimensional assay with low serum and low cell numbers, the D2.OR cells remained dormant whereas the D2A1 cells proliferated^29^. Several slightly adapted 3-dimensional culture assays have since been used to study the D2 cells *in vitro*, which all recapitulated the *in vivo* findings^21,25^. Here, we made use of the Matrigel on Top assay (MoT)^21^. Consistent with previous reports, we could show that both the D2.OR and the D2A1 cells proliferate equally in 2D, whereas the D2.OR cells hardly expanded, but the D2A1 cells did expand, in the 3D MoT assay (**Fig 2a and Supplementary Fig 4a**). To assess if the lack in growth expansion in the D2.OR cell in 3D was caused by a growth arrest, or because of a balance between proliferation and death, we determined the percentage of mKi67+ cells and the amount of cleaved caspase-3 (CC3) in the cells in 2D and 3D. Overall, the D2A1 cell line had a slightly higher CC3 score compared to the D2.OR (**Supplementary Fig 4b**). No increase in CC3 staining was observed for the D2.OR cells in 3D when compared to 2D. None of the D2.OR cells in 3D were positive for KI67, in contrast to the cells in 2D and to the D2A1 cells in either growth condition (**Supplementary Fig 4c**). These data suggested that a cell cycle arrest was underlying the growth reduction in the D2.OR cells in 3D. To corroborate the D2.OR cell cycle arrest, we loaded dormant D2.OR cells with a DiI and studied its retention or dilution over multiple days. Indeed, DiI was retained in dormant D2.OR compared to the D2A1 cells in 3D as determined by FACS and (live cell) microscopy (**Fig 2b and supplementary Fig 4d-f**).

**Fig. 2:**
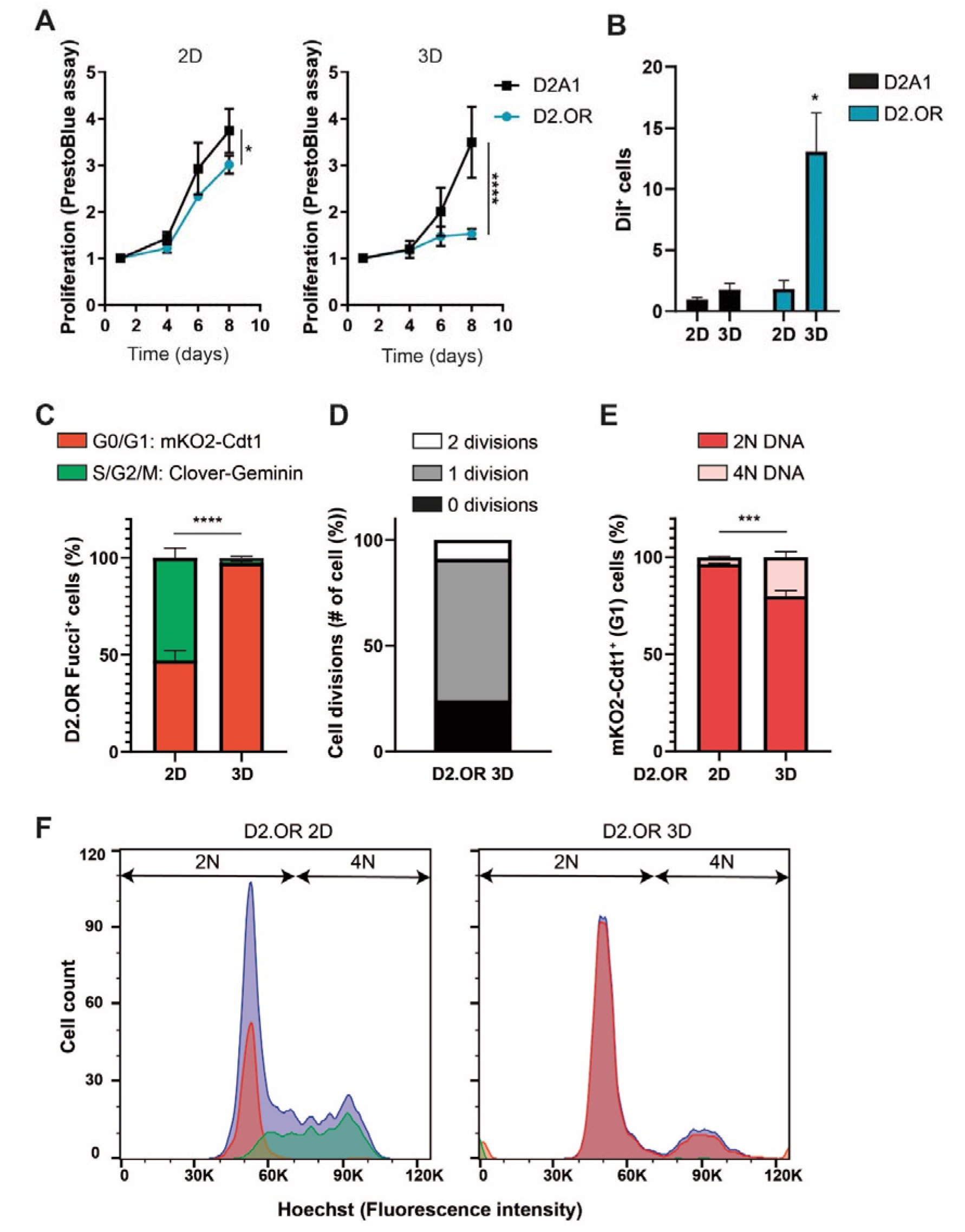
Characterization of D2.OR G1 cell cycle arrest in 3D culture. **a** Proliferation analysis of D2A1 and D2.OR cells in 2D or 3D culture conditions using PrestoBlue cell viability assay. N = 3 replicates in duplo/triplo. 2D graph: Interaction effect, *F* (3,42) = 3.685, P = *. 3D Graph: Interaction effect, *F* (3,42) = 10.57, P = ****. **b** Quantification of the percentage of DiI^+^ cells after 8 days of culture by FACS. N = 3 repeats. Interaction effect, *F*(1,6) = 6.278, P = *; *post hoc* test, D2OR 3D vs all others P = *. **c** Quantification of the percentage of Fucci^+^ cells (either Clover^+^ or mKO2^+^) after 3 days of culture by FACS. N = 3 repeats. Interaction effect: F (1,10) = 256.3, P = ****; *post hoc* test, 2D vs 3D mKO2 P = ****; **d** Quantification of the number of cell divisions until cells remained in G1 (mKO2+) after plating in Matrigel. Cells were imaged every 3 hours. N = 5 positions, 13 cells. **e** Quantification of FACS analysis showing the percentage of mKO2^+^ D2.OR Fucci+ cells containing 2N or 4N DNA (measured by Hoechst) in 2D or 3D after 3 days of culture. N = 3. Interaction effect: F (1,10) = 46.11. P = ****. *post hoc* test, 2D vs 3D 4N P = ***. **f** FACS plot of experiment quantified in c and e. Blue: Hoechst, Green: Clover-Geminin, Red: mKO2-Cdt1. P values were calculated using a (repeated measures) 2-Way ANOVA.

Dormancy is often defined by a G0/G1 arrest. We made use of the FUCCI cell cycle fluorescent reporter to determine the dormant D2.OR cell cycle status^35^. We engineered D2.OR to stably express fluorescently-labelled cell cycle regulators chromatin licensing and DNA replication factor 1 (Cdt1)_30-120_ and Geminin_1-110_. mKO2-Cdt1_30-120_ is expressed in G1 (red color), both Cdt1-mKO2 and Clover-Geminin_1-110_ are expressed in S (yellow color), and Clover-Geminin1.110 is expressed in G2/M phase (green color). After validating the FUCCI reporter system (**Supplementary Fig 5a**) we assessed the cell cycle status of D2.OR_Fucci cells *in vitro* by fluorescence activated cell sorting (FACS) and microscopy. During 2D proliferation, between 35-45% D2.OR cells were in G0/G1, whereas in the MoT, >99% cells were arrested in G0/G1 (**Fig 2c and supplementary Fig 5b**). To assess when and if cells divide after plating in MoT, we performed live-cell imaging over 69 hrs. Interestingly, we observed that immediately after plating, D2.OR cells demonstrated a migratory and stretched phenotype (**Supplementary Fig 5c**). When cells made new contacts, they clustered together and thereafter remained in this cluster (**Supplementary Fig 5c and Movie 1**). At the end of the observation period, cells showed a rounded phenotype (**Supplementary Fig 5c and Movie 1**). We furthermore observed that most cells underwent 1 cell division, although some underwent no or 2 cell divisions (**Fig 2d**). The median time it took before all cells in a cluster were in G0/G1 was 24 hours and by day 3 all cells were in G0/G1 (**Supplementary Fig 5d**). Altogether, these results confirmed the G0/G1 cell cycle arrest of the D2.OR cells in 3D, and suggest that cells undergo a maximum of two cell cycles after plating in Matrigel.

### Dormant D2.OR can be 4NG1

G1 cells are normally diploid (2N), however, it has been observed that some G1 arrested cells are tetraploid (4NG1). This G1 arrest is important to prevent tetraploid cells becoming aneuploid^36,37^. We thus wondered what the ploidy of dormant D2.OR cells was. We first performed karyotyping, and concluded that both cell lines are polyploid (**Supplementary Fig 6a**). But where D2.OR cells are mostly triploid, D2A1 cells are mostly hypotetraploid. Next, we performed a Hoechst FACS cell cycle analysis on dormant D2.OR_Fucci cells plated in 3D. For simplicity, we refer to cells that have not replicated their chromosomes yet in S phase to 2N, despite them being polypoid (**Supplementary Fig 6a**) and thus having three or sometimes four copies of the same chromosome. As expected, in 2D, the G1 cell cycle status matches the 2N state, and the G2 cell cycle status matches the 4N state (**Fig 2e-f and Supplementary Fig 6b**). Interestingly, in 3D, a significant population (~20%) of G1 cells is 4N (**Fig 2e-f and Supplementary Fig 6b**). Thus, some dormant D2.OR cells reside in 4NG1.

### Dormant D2.OR genotype does not correlate with senescence signature

Dormancy and senescence are both characterized by cell cycle exit, and dormant-like senescence programs have been postulated^3,15^. As such, we wondered if the D2.OR cells in the MoT model should be considered dormant or senescent. To assess this in an unbiased manner, we performed RNA-seq on the D2.OR and D2A1 cells in 2D and 3D (**Fig 3a**) after 7 days of culture. We aimed to identify those genes that showed a significant interaction between the cell lines and the culture type. Once we generated a ranked list of genes based on their statistical significance, we performed a GSEA analysis to identify the enriched pathways at both end of the list. Gene sets related to proliferation and cell cycle (mitosis) showed a negative normalized enrichment score (NES), i.e. higher log_2_ fold change (3D - 2D) in D2A1 than in D2.OR (**Fig 3b**). Interestingly, similar to our previous study in an unrelated dormancy model, we identified gene sets that associated with the dormancy D2.OR phenotype (i.e., higher log_2_ fold change (3D - 2D) in D2.OR than in D2A1) were related to lipid and alcohol metabolism^36^.

**Fig. 3:**
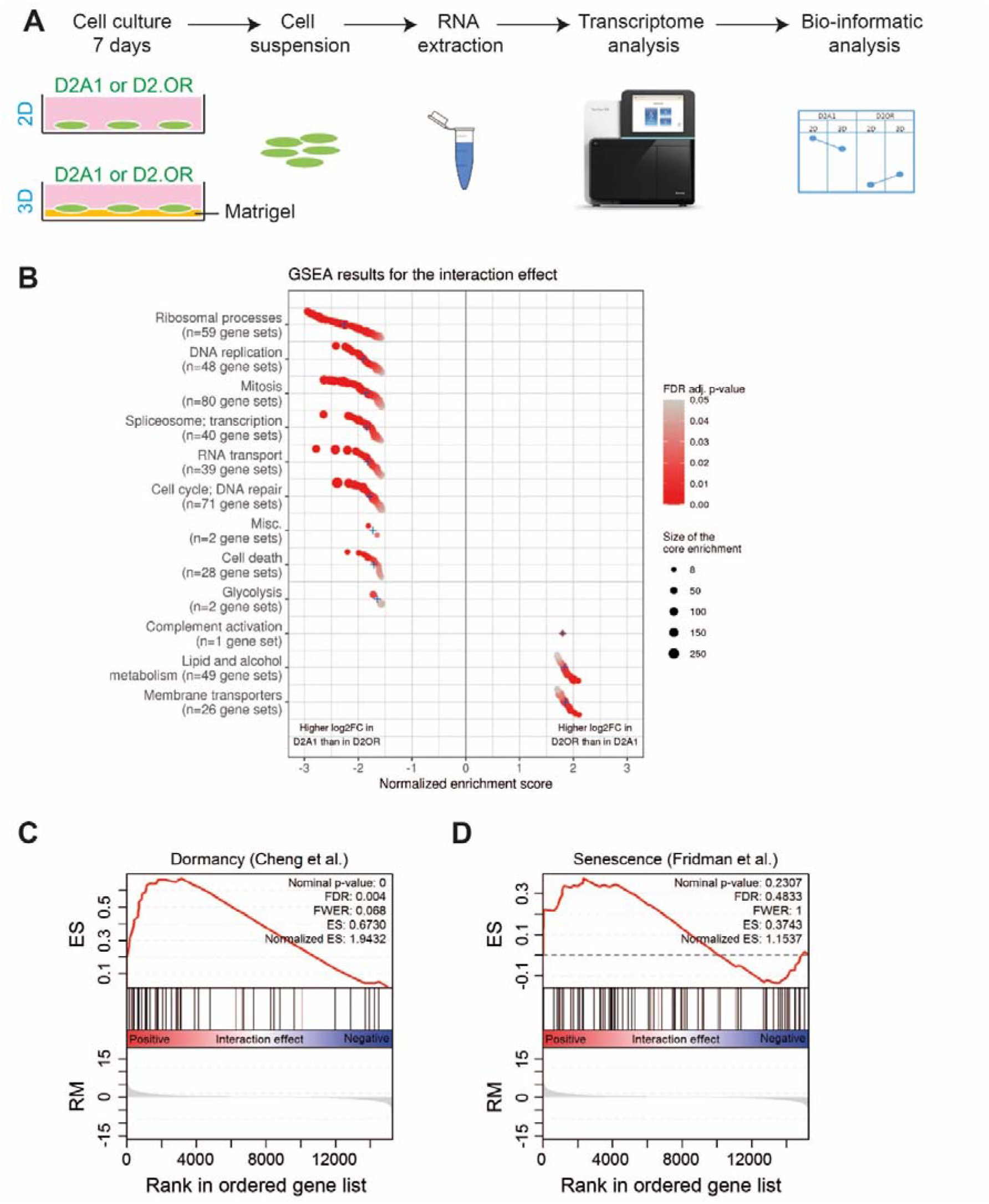
Unbiased transcriptomic analysis suggests a D2.OR dormant cell cycle arrest. **a** Schematic representation of the experimental pipeline. **b** GSEA analysis on the interaction effect when comparing D2A1 and D2.OR in 2D and 3D. Gene-sets ordered by normalized enrichment score (NES) within each cluster. Blue crosses represent average NES in each cluster. Point color represents FDR-adjusted p-value and size represents core enrichment. Log2FC = log2 fold change. **c-d** GSEA analysis of a published dormancy signature (Cheng *et al* ^54^) (**c**) or senescence signature (Fridman *et al* ^55^) (**d**) on the interaction effect when comparing D2A1 and D2.OR in 2D and 3D. ES, Enrichment score; RM, ranking metric.

Next, we performed pre-ranked GSEA analyses of published dormancy and senescence gene sets using the results of the interaction analysis as the ranking metric. The dormancy signature by Cheng *et al* was found to be positively enriched (FDR < 0.05) in the D2.OR 3D cells (i.e., higher log_2_ fold change (3D - 2D) in D2.OR than in D2A1) (**Fig 3c**). Additionally, we did not observe a significant enrichment for the senescence signature (**Fig 3d**).

Thus, gene sets associated with cell cycle progression were negatively enriched, and dormancy-related gene sets were enriched in our analysis, pointing more towards a dormancy phenotype for the D2.OR cells in 3D. A complete list of gene sets tested and their results is available in Supplementary table 2.

### Pathways involved in senescence are not active in dormant D2.OR

Molecularly, dormancy is a different process from senescence. Senescence is not defined by specific markers, but rather by a multi-marker approach^37^. To determine if a senescence program might be responsible for the observed cell cycle arrest, we assessed the following senescence associated markers: SA-β-galactosidase, DNA damage (γ-H2AX), and the loss of Lamin B1.

We observed that dormant D2.OR expressed β-galactosidase as much as the senescence control (MCF7 treated with etoposide in 2D), whereas the proliferative control (D2A1) did not (**Fig 4a**). In contrast, Lamin B1 loss was not observed in dormant D2.OR, whereas it was lost in a senescence control (irradiated MCF7 cells) (**Fig 4b and Supplementary Fig 7a**). Finally, senescent cells are known to accumulate DNA damage. We therefore checked DNA damage through γ-H2AX staining. Upon DNA double-stranded breaks, γ-H2AX gets phosphorylated and accumulates in the cell forming small foci. Interestingly, we did not find an increase of foci accumulation in dormant D2.OR compared to control (D2.OR 2D), and overall the intensity was much less compared to a senescence control (HeLa cells treated with etoposide) (**Fig 4c and Supplementary Fig 7b**). As two out of the three senescence markers were not present in our phenotype, this argues against senescence.

**Fig. 4:**
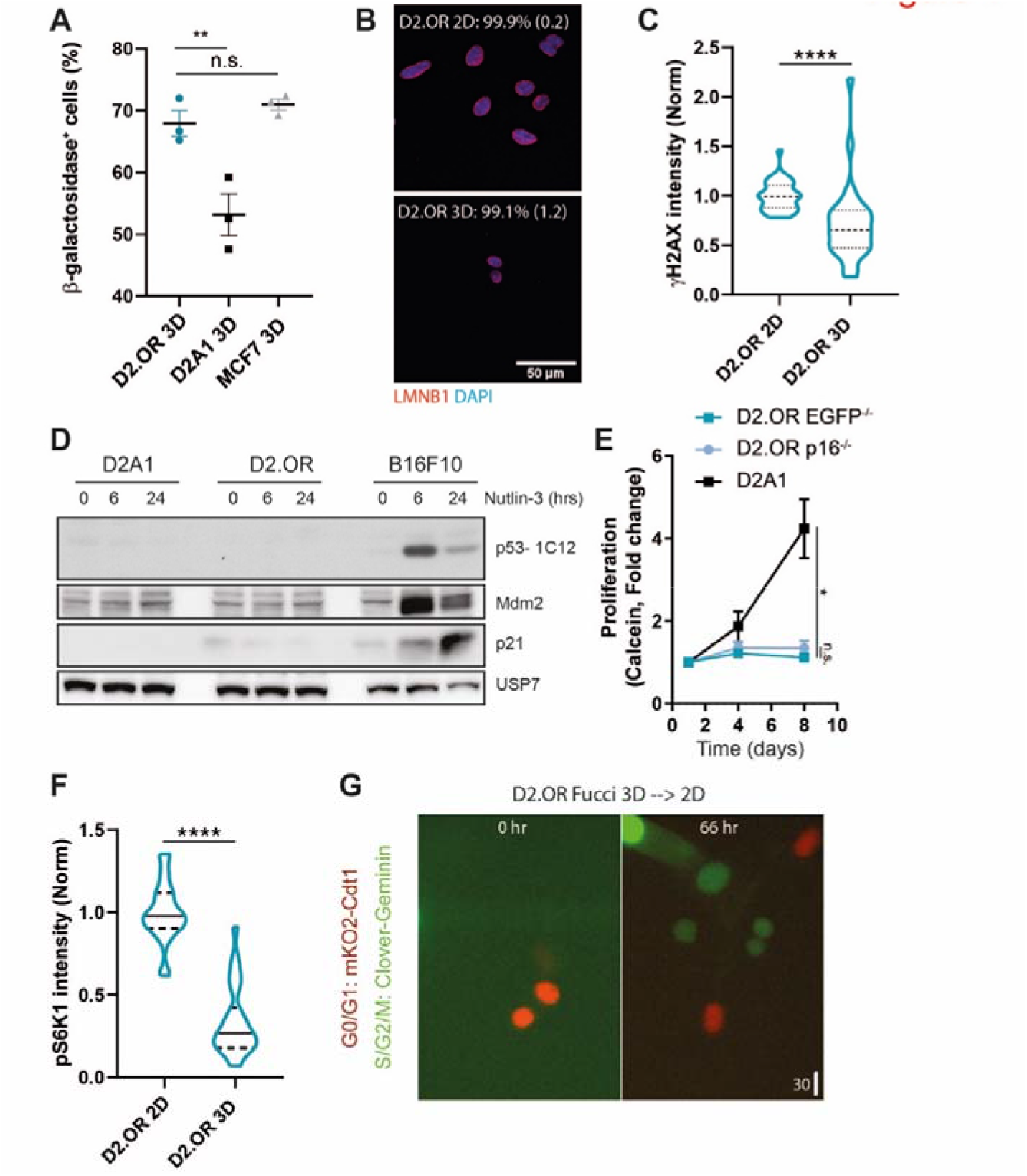
D2.OR cell cycle arrest in 3D is associated with dormancy and not senescence. **a** Quantification of the percentage of β-galactosidase+ cells cultured for 4 days in 3D. MCF7 was used as a senescence control. N = 3 replicates. **b** Microscopy images of D2.OR cells cultured for 4 days in 2D or 3D conditions, stained for DAPI and LaminB1. Percentage of positive laminB1 cells (±SD) is indicated. N = 3 replicates, each ≥ 12 cells. **c** Quantification of nuclear γH2AX staining intensity of D2.OR cells cultured for 4 days. Replicates normalized to 2D. N = 30 field of view (FOV) of 3 replicates. **d** Immunoblot analysis of lysates from indicated cell lines cultured in 2D and treated with Nutlin-3 [10 μM] for 0, 6 or 24 hours. **e** Quantification of proliferation of indicated cell lines cultured for 8 days in 3D. Replicates were normalized to day 1. N = 3 replicates, in duplo. Time x Cell line, F (4, 27) = 10.68. P = ****. *Post hoc* test, day 8, D2A1 vs others P = *, D2.OR EGFP vs P16-/- P = n.s. **f** Quantification of pS6K1 immunostaining of D2.OR cells cultured in 2D or 3D. Replicates are normalized to 2D. N ≥ 24 field of view (FOV) of 3 replicates. **g** Microscopy images of a time series of D2.OR Fucci cells in 2D. Cells were cultured in 3D for 4 days, extracted, re-plated in 2D, and imaged. P values were calculated using 1-way ANOVA, Mann-Whitney test or mixed effects analysis.

p53 and p16 are the main pathways that can induce senescence, thus we wanted to investigate if p53 and/or p16 would be required for the observed cell cycle exit in D2.OR cells in 3D. When we assessed the response of both cell lines to Nutlin-3, which stabilizes p53 protein levels resulting in an increase in downstream target gene expression, neither of the D2 cell lines showed an increase in mRNA levels of the two p53 target genes, p21 and MDM2, while these genes were increased in a positive control cell line (B16F10) (**Supplementary Fig 7c**). Similar results were observed at the protein level (**Fig 4d**). Moreover, we were unable to detect p53 protein in either the D2.OR or D2A1 cell line (**Fig 4d**). Combined, the data suggest that neither the D2.OR cells nor the D2A1 cells contain functional p53. P16, however, was expressed by D2.OR cells, but decreased in D2.OR cells in 3D (**Supplementary Fig 7d**). In addition, CRISPR-Cas9-mediated knockout of the *p16* gene did not result in escape from dormancy in the MoT assay (**Fig 4e and Supplementary Fig 7e**). These results indicated that the two main senescence-inducing pathways p53 and p16 were not responsible for the D2.OR cell cycle arrest phenotype.

Next, we tested if dormancy markers were associated with the observed cell cycle arrest. Cell cycle inhibitor p27 is often associated with dormancy, whereas p21 and p16 are mostly associated with senescence. p27, but not p21 or p16, showed a specific increase in the D2.OR cell line and not the D2A1 line when comparing 2D versus 3D (**Supplementary Fig 7d**), suggesting a dormancy program. Dormancy is often associated with low levels of mammalian target of rapamycin (mTOR) (e.g. due to low nutrient levels), whereas during senescence, mTOR levels are often high ^37–39^. We therefore assessed the activity of mTOR by performing immunofluorescence of downstream kinase target S6K1. Phosphorylation of S6K1 suggests higher mTOR activity. D2.OR cells plated in 3D had significantly lower p-S6K1 levels compared to D2.OR cells plated in 2D. As expected, senescent cells (Hela cells treated with etoposide) did not show a reduction in p-S6K1 (**Fig 4f and supplementary Fig 7f**). Another hallmark for dormancy is the reversibility of the cell cycle arrest. And even though reversible senescence has been observed, it is not that common, and it has only been observed for cells that have just initiated a senescent program^40^. Thus, we assessed if arrested D2.OR were able to escape from dormancy and start proliferating (assessed by Fucci, or by PrestoBlue cell viability assay) when cells were recovered from Matrigel and re-plated in 2D. Indeed, we observed that the dormancy phenotype was reversible (**Fig 4g, Supplementary Fig 7g and Movie 2**). Combined, we conclude that the cell cycle exit phenotype of D2.OR cells in 3D is not associated with a senescence phenotype, but rather with a reversible dormant phenotype.

## DISCUSSION

Dormancy is incompletely understood, but its importance in cancer progression and resistance is evident as reactivation of dormant cancer cells can lead to cancer relapse and eventually patient death.^6,7^ Proper dormancy models and a thorough understanding of these models are essential to expose the molecular and cellular characteristics of dormancy. Such models should be specific for dormancy, and should not model other cell cycle arrest processes like senescence. Here, we evaluated the widely used D2 breast cancer cell lines model for its capacity to specifically recapitulate dormancy *in vitro* and *in vivo*.

Similar to others we were able to show that the D2.OR cell line underwent a cell cycle arrest in an experimental metastasis model. However, we are the first to show that D2.OR become dormant in the liver of syngeneic mice, i.e. mice with a fully competent immune system. This is important as the adaptive immune system has been shown to regulate single cell dormancy^25^. Moreover, by combining label retention and immunohistochemistry, we were able to specifically address proliferation and death in the truly dormant cell population. Indeed, all label retaining cells were negative for KI67 and cleaved caspase 3, excluding “balanced dormancy” which is based on equal number of proliferating and dying cells^11^. Interesting is that not all D2.OR cells remain dormant in the liver over the course of two weeks. This heterogeneity may be caused by the microenvironment, e.g. through variations in stiffness, or alternatively by cellular heterogeneity. The latter is emphasized by the genomic instability shown as variation in chromosomal numbers and rearrangements we observed between the different metaphase cell derived nuclei from the same cell line.

Dormancy is linked to tumor recurrence and worse prognosis. Recurrence is more likely to occur in ER^+^ and HER2^+^ patients than in triple negative patients. Similarly, recurrence is more associated with Luminal A and B subtype than the basal subtype. Surprisingly, we determined that the D2 cells most likely belong to the ER^+^/Her2^+^ basal subtype. This correlates with a small proportion of patients, which are at high risk for disease recurrence, but which show significantly improved response to neoadjuvant chemotherapy.^41^ These patients are ER^+^, but show low levels of ER at the transcriptional level, which was attributed to high expression of dominant negative ERΔ7 mRNA. Indeed, we also show that the cells have low expression of ER mRNA, but they are clearly responsive to 4OHT, suggesting they should have ERα at the protein level. Indeed, Raj *et al* show that D2A1 cells have ERα^42^.

Our results are in contrast to a recent report from Yang *et al*, who classified D2A1 cells as being Luminal B, based on a clustering analysis of transcriptomic data^43^. However, depending on the computational model they used, D2A1 cells were assigned to different subclasses (2x basal, 2x LumA, 2x LumB and 2x Her2). Interestingly, when Genefu and Support Vector Machine methods were used, D2A1 cells were assigned to the basal category^43^. This indicates D2A1 cells are not so easily classified, and might explain the confusion in the literature. As such, it might be best to define the cells based on hormone receptor presence, ER+/Her2+. Overall, this suggests the D2 cells are excellent cell lines to model ER+/Her2+ disease recurrence.

The identification of 4NG1 D2.OR cells when cultured in 3D is surprising. So far, this has never been associated with cells in dormancy, but rather with senescence.^44^ 4NG1 cells can appear if cells are delayed in mitosis (D-mitosis). In the presence of an active spindle assembly checkpoint, cells in D-mitosis can ultimately escape (termed mitotic adaptation or slippage) and enter the next G1 as tetraploid cells.^44^ These 4NG1 are either arrested in senescence, or are able to proliferate normally. We, on the other hand, suggest that the 4NG1 D2.OR cells cultured in 3D are in dormancy. It will be of interest for future research to determine the fate of the 4NG1 cells. E.g. are these cells also able to escape dormancy and resume proliferation, or have these cells in fact entered a permanent senescence state.^44^ It is important to note that we cannot exclude the latter; the overall population of D2.OR cells in 3D is in dormancy, but a small proportion (e.g. the ~20% 4NG1 cells) might be in senescence. In addition, it will be important to determine if the 4NG1 cells are present *in vivo* and in other model systems as well. At last, it will be of interest for future research to determine the reason for the 4NG1 arrest. Studies have shown that absence of p53 protein can accelerate the exit from D-mitosis.^45^ As the D2.OR cells are p53 negative, this might explain the presence of 4NG1 cells. However, why the cells were in D-mitosis is unclear, as this is normally induced by treatments that depress or altered microtubule assembly/dynamics (e.g. aurora-kinase inhibitors).^45^ We showed that the D2.OR cells are undergoing a dormant cell cycle arrest. Why do some of the arrested D2.OR cells in 3D show increased expression of β-galactosidase? It is important to note that none of the senescence markers are specific, and are in fact also expressed by other cells under certain circumstances. Indeed, β-galactosidase expression has been reported in other dormant cells.^13^ Thus, senescence can only be confirmed when a multitude of markers is present,^37^ which was not the case for the D2.OR cells in 3D.

Instead, we confirm that the D2.OR cells in 3D are in a dormancy arrest. Based on transcriptomic analyses, we show that this phenotype is associated with gene sets related to lipid and alcohol metabolism. When dormant cells in a completely unrelated 2D assay were assessed, a similar association was found, and this was linked to exosome secretion by dormant cells.^36^ This suggests that exosome secretion might be a general characteristic of dormant cells, but it has to be confirmed in the D2.OR model system.

The discrimination between a senescence or dormancy cell cycle arrest program is important when considering treatment options. As the pathways that define either program are quite distinct^39^, targeted treatment for one program will most likely not work against the other program. For example, FOXO4-DRI senolytic eliminates senescent cells by inhibiting the interaction between FOXO4 and p53. As p53 is not expressed in D2.OR dormant cells, this senolytic will most likely not eliminate these cells. Which drugs will eliminate dormant cells is subject of active investigation, and can be pursued with the D2.OR dormant model system.

## METHODS

### Cell culture

Mouse mammary carcinoma cell lines D2A1 and D2.OR were obtained from Karmanos Cancer Institute (F.R. Miller)^16^. Mammary cancer cells D2.OR and D2A1, mouse embryonic fibroblast (MEF) cells, mouse melanoma cancer cells B16F10, human breast cancer cells MCF-7, human cancer cells Hela, and human kidney cells HEK293T were all cultured in Dulbecco’s Modified Eagle Medium (DMEM) high glucose (Gibco), supplemented with 10% foetal calf serum (FBS) (biowest), 60 μg/mL penicillin (Roth) and 96 μg/mL streptomycin (Roth). Cells were maintained in a 5% CO2 humidified incubator (Forma Scientific) at 37 °C. Every month, all cell lines in culture were subjected to a mycoplasma infection test.

For the standard two-dimensional (2D) experiments, cell density was determined using the TC20™ Automated Cell Counter (Bio-Rad). Thereafter, 1×10^3^ cells were seeded per well in either a 48-well or 8-chamber format in DMEM high glucose (Gibco) and refreshed every 3 to 4 days.

The Fucci reporter construct (pLL3.7m-Clover-Geminin(1-110)-IRES-mKO2-Cdt(30-120)) was a gift from Michael Lin (Addgene plasmid #83841)^35^. Stable Fucci reporter expression and H2B-Dendra2 expression was performed by lentivral particles transduction. Transfection of HEK293T cells by polyethylenimine (PEI) with a viral vector and helper constructs was performed in 10 cm culture dishes. 6mL of virus containing medium was filtered and supplemented with 10 μg/mL polybrene. The suspension was used for the infection. pLV_CMV_H2B-dendra2 cell selection was performed with medium containing 1.5 μg/mL puromycin, whereas the selection of Fucci was performed by flow cytometry.

### Matrigel-on-top assay

For the Matrigel on top culture assay^46^, 48-wells plate were coated with 40uL/Well of (growth factor reduced) matrigel (Corning) and incubated at 37°C for 30min, which allowed the Matrigel to solidify. Subsequently, cells (1000 cells/well) were resuspended as single cells in 200uL of 3D media (Nutrient Mixture F-12 (DMEM/F-12) + GlutaMAXTM-I (Gibco) supplemented with 2% Horse serum (Gibco), 0.5 μg/mL hydrocortisone (Sigma Aldrich, H4001), 50ng/mL cholera toxin (Sigma Aldrich, C8052), 10 μg/mL insulin (Sigma Aldrich, I3536), 60 μg/mL penicillin (Roth), 96 μg/mL streptomycin (Roth)) containing 2% Matrigel, and added on top of the solidified Matrigel. For the 2D culture, cells were seeded directly on plastic in the 3D media. Medium was refreshed every 3 to 4 days.

### Cell viability assay

To determine the proliferation rate in 2D or 3D, cells were cultured for ≤8 days in 48-wells plate as described in the cell culture section. At desired timepoints, 20 μL of PrestoBlue cell viability reagent (Invitrogen) was added directly to the media and the plate was incubated for 10 minutes at 37°C in the dark. Subsequently, fluorescence at wavelength 544/590 nm was captured using the VICTOR X3 plate reader (PerkinElmer). Measurements were normalized to controls with only Matrigel and/or media to adjust for background signals.

Calcein-AM was additionally used to determine proliferation^47^. In short, cells were washed twice with Phosphate-Buffed Saline (PBS) after which 5 μM calcein-AM (Thermofisher, C1430) in PBS was added. After incubating for 30 minutes at 37 °C in the dark, fluorescence was captured at wavelength 494/517 nm using the VICTOR X3 plate reader (PerkinElmer). Afterwards, measurements were normalized to controls with only Matrigel and/or media to adjust for background signals.

### DiI loading

Cells cultured in 2D were trypsinized and counted. 1×10^6^ cells/mL were incubated for 7min with 5uL/mL DiI vibrant solution. Loaded cells were spin down 3min at 1300rpm and pellet was washed with PBS 2 times. Cells were counted again before plating in 2D or 3D condition. For animal experiment, cells were always loaded with DiI one day prior to the injection.

### FACS

For the 2D condition, D2.OR-FUCCI4 cells were collected and diluted to 1×10^6^ cells/mL suspension in 0.2% BSA in PBS. Thereafter, 2×10^6^ cells were incubated with Hoechst 33342 (1 mg/mL) (Thermofisher Scientific, 62249) to stain the nuclei. After incubation in the dark for 30 minutes at 37°C, cells were collected in 500 μL PBS and subjected to flow cytometry.

For the 3D condition, D2.OR-FUCCI4 reporter cells were seeded in 6 cm dishes with Matrigel, cultured for 4 days and directly stained with Hoechst 33342 (1 mg/mL) (Thermofisher Scientific, 62249) for 30 minutes at 37°C. Thereafter, cells were extracted from Matrigel according to our self-optimized protocol in which every step is performed on ice and requires all tips to be pre-coated with 2,5% BSA (Sigma Aldrich) in PBS to yield the highest number of living cells. The cells were washed two times with PBS, ice-cold Cell Recovery Solution (Corning) was added and cells were gently rocked to facilitate for 20 to 30 minutes on ice in order to dissolve the Matrigel. After dissolvement was confirmed by microscopy, 1% BSA (Sigma Aldrich) was added to the cell suspension and cells were centrifuged twice (approximately 280g at 4°C) for 5 minutes. After discarding the supernatant, the cells were resuspended in 500μL 0.5% BSA (Sigma Aldrich) in PBS and subjected to flow cytometry.

Flow cytometry was performed using the BD LSR II machine and BD FACSDiva software. Afterwards, recordings were analysed using the FlowJo software.

### Microscopy

Confocal imaging was conducted on the Leica SP5. Images were acquired as sequential scans and collected in 8- or 12-bit. Plastic 48-well plates were imaged using the dry 40x long distance magnification, whereas the glass 8-well chamber slides were imaged using the 63x oil objectives. The fluorescent detection range was set manually based on the emission spectra of the different fluorophores. We used the following lasers depending on the fluorophores: 405 Diode, Argon, DPSS561, HeNe594 and HeNe633. Furthermore, Z-stacks differed between 1,5 μm and 3,0 μm depending on the protein of interest.

Live-cell imaging was conducted on the Leica AF6000 and Leica DMI6000. For both, plastic 48-well plates were imaged using the dry HPX PL FLUOTAR L 40×0.6NA long distance objective. The following filters cubes were used: geminin-Clover, Triple filter G; cdt1-mk02, RFP and Brightfield: empty. Live-cell imaging experiments with FUCCI4 expressing D2.OR and D2A1 cells were performed for 1 to 4 days with frames every 20 minutes to 3 hours. Z-stacks were adjusted for individual positions and varied between 2 and 3 μm. Live cell imaging of D2.OR and D2.A1 cells loaded with DiI were performed for 93 hours with frames every 30 minutes. Furthermore, temperature and CO2 percentage, at respectively 37°C and 5%, were maintained using an environmental control system.

Images were gathered with confocal and live-cell imaging analysed using ImageJ software.

### Immunofluorescence

Cells were cultured on 48-well plates (CELLSTAR), μ-Slide 8 Well (Ibidi) or Nunc Lab-Tek chamber slides (ThermoFisher Scientific). Cells were fixed in 2% paraformaldehyde for 10min and permeabilized in PBS supplemented with 0.5% Triton X-100 for 10 min at room temperature (RT). Then cells were washed 2 times in PBS supplemented with 100mM Glycine (PBS glycine) for 10 min and incubated with blocking buffer (130mM NaCl, 7,7 mM NaN3, 0.1% bovine serum albumin, 0.2% Triton ×100, 0.05% Tween 20, 5% goat serum) for 1h at RT. Primary antibody were diluted in blocking buffer at the following concentration: Ki67 (ab16667, AbCam, 1:200), Cleaved-Caspase3 (9661, cell signaling, 1:200), Anti-phospho-Histone H2A.X (Ser139) (clone JBW301, 05-636 Millipore, 1:1000), p-p70 S6 kinas a (pS6K1)(Santa cruz, sc-8416, 1:100), LMNB1 (abcam, ab16048, 1:500) and incubated overnight at 4°C. Subsequently, cells were washed 2 times 10 min with PBS Glycine and incubated with A488 conjugated secondary antibody diluted in blocking buffer at 1/200 for 1h at RT. Finally, cells were mounted in media containing DAPI (Hard vectashield set) and immediately analyzed using the Leica SP5 confocal microscope or stored at −20C to prevent Matrigel dissolution and fluorescent signal degradation.

### Immunoblotting

Cells were lysed in Laemmli buffer (12% 1M Tris-Cl pH 6.8, 4% SDS, 20% glycerol, ddH2O) supplemented with 1x phosphatase inhibitor cocktail (P5726, Sigma), and total protein concentration was determined using the DC protein assay (Bio-Rad). In total, 10 μg of protein was loaded per lane. The proteins were separated by 10% sodium dodecyl sulfate polyacrylamide gel electrophoresis and transferred onto a polyvinylidene difluoride membrane using electrophoresis for 1 hour. Following, the membrane was blocked at room temperature for 1 hour in tris-buffered saline containing Tween-20 (TBS-T) containing 5% milk powder. The membrane was washed with TBS-T and subsequently incubated with the primary antibodies at 4°C overnight. The types of primary antibodies used were: Anti-CDKN2A/p16INK4a ([EPR20418] (Abcam, ab211542)), Anti-Vinculin (hVIN-1 (Sigma Aldrich, V9131, 1:1000)), anti-USP7 (Bethyl Laboratories, A300-033A), anti-MDM2, clone 3G9 (Merck Millipore, 04-1530), anti-p53 clone 1C12 (Cell Signaling Technology), antip21, Clone F5 (Santa Cruz Biotechnology). After washing with TBS-T followed with incubation with horseradish peroxidase (HRP)-conjugated anti-rabbit, anti-goat, or anti-mouse secondary antibody (Amersham science, 1:10.000) at room temperature for 2 hours. Bands were detected using enhanced chemiluminescence (Clarity ECL western blotting substrate, 1705061, Bio-Rad) in accordance with the supplier’s protocol.

### qPCR

Total RNA of cells was extracted using the Macherey-Nagel RNA kit (Thermo Fisher Scientific, Netherlands). cDNA was made from 1000 ng RNA mixed with 5x reaction buffer, Rnase inhibitor and 10 mM dNTP (ThermoFisher, Netherlands) using the Thermo Fisher Scientific PCR protocol, 5 min at 25°C, 60 min at 42°C, 5 min at 70°C. PCR was performed on the T100 Thermal Cycler (Bio-Rad Laboratories, USA). cDNA was diluted to 100 ng/μL. A mastermix of RNase free mQ, forward primer and reverse primers was added to cDNA along with SybrGreen (ThermoFisher, USA). Gene expression was measured with the BioRad CFX Man 3 (Bio-Rad laboratories, CA, USA). The measured gene expression was corrected with housekeeping gene *Gapdh*, or with the geomean of *Hmbs* and *Rpl13a*.

### β-galactosidase assay

Cells were seeded 1×10^3^ cell/well in a μ-Slide 8 Well (Ibidi, 80826) chambered coverslip in both 2D and 3D. After culturing for 4 days, cells were washed twice with PBS. Thereafter, the β-galactosidase assay was performed using the Senescence beta-Galactosidase Staining Kit (Cell Signaling, 9860S) according to the supplied protocol.

Cells were fixed with 1X fixative solution for 15 minutes at RT, washed with PBS and stained with 1X β-galactosidase staining solution at pH 6.0. After staining, the chambered coverslip (Ibidi) was stored in a culture dish (Cellstar), sealed with parafilm and incubated in a dry incubator at 37 °C in the absence of CO2. After 3 hours of incubation cells were imaged in brightfield using the Leica DMi8 microscope with 20X magnification. Images were quantified manually by counting the percentage of positive clusters.

### Irradiation

MCF-7 breast cancer cells were γ-irradiated with a total of amount of 10 gray (Gy) (1 Gy/min) using the YXLON X-ray apparatus (200 kv/4,0 mA – foc = 5,5). Lead plates on top of the chamber coverslips (Ibidi; 80826) were used to protect other cells from γ-radiation. After irradiation, the cells were cultured under normal circumstances for subsequently 24, 48, 72 or 96 hours before they were fixed and stained for immunofluorescence.

### Reversible growth arrest assay

D2.OR Fucci cells were cultured as described in the Matrigel-on-top section. Instead of a 48 well plate, the culture was upscaled to a 6 well plate. Cells were extracted from the Matrigel as described in the FACS section. After extraction, cells were plated in 2D in a μ-Slide 8 well (ibidi, 80826), and imaged on a Leica AF6000 LX with a Hamamatsu-C9100-02-COM4 camera and a HCX PL APO CS 20.0×0.75 DRY UV objective in 14 bit every 20 minute for 3 days. Multiple positions were recorded and analysed by ImageJ.

Alternatively, D2OR cells were cultured in the Matrigel on Top assay as described in the Matrigel-on-top section. 7 days after seeding, cells were extracted from Matrigel. Media was removed from the well and 200 μL of ice cold trypsin was added. Mechanic disruption of the Matrigel was performed using P1000 tips and cycle of pipetting up and down. The suspension was then transfer to a 15 mL tube and incubated for 5 min at 37°C. Again mechanic disruption by pipetting up and down was performed using a P1000 and 10mL of ice cold PBS was added. Cells were centrifuged for 3 min at 1300rpm and supernatant was discard. A second wash with PBS was performed before resuspending the pellet in 3D media. 16 wells from a 48-wells plate were pooled to re-seed 1000 cells/well in 2D and 3D. Seeding was performed as described earlier in this section. Proliferation readout using Calcein-AM (see section Proliferation assay) was performed at day1 and day7 to determine the dormant/proliferative phenotype of the re-seeded cells.

### Label retention

Vybrant DiI cell-labeling solution (5 μl/mL, V22888, Invitrogen) was directly added to cell suspensions at a density of 1×10^6^ cells/mL, and incubated for 7 minutes, followed by thorough rinsing with phosphate buffered saline (PBS). Cells were seeded at 1×10^3^/well in a 48-well plate, and retention of the label was determined after 8 days in culture using fluorescence-activated cell sorting (FACS) or microscopy. FACS experiments were carried out on a BD LSR IIFlowcytometer (BD Biosciences), and subsequently analysed using BD FACSDiva software (BD Biosciences). To determine the label retention for each condition, cells were harvested after 8 days of culture, washed, and analysed by flow cytometry. Cells cultured in 3D were harvested from the basement membrane matrix using Cell Recovery solution (Corning), in accordance with the manufacturer’s instructions. Cell morphology was observed by light microscopy, time lapse images were captured by an AF6000 inverted widefield microscope (Leica Microsystems) and label retention was quantified using confocal laser scanning microscopy, for which the Leica SP5 was used with a 40x long working distance objective, after which images were processed and analysed using ImageJ Fiji software

### Transcriptomic analysis

D2OR and D2A1 were seeded as described in the Cell Culture section. 10 wells of each cell line were pooled to be able to extract enough RNA. Cells were extracted from the Matrigel using the recovery solution from Corning. Briefly, media was removed and cells were washed 3 times with PBS. 140uL of ice cold recovery solution was added per well. After 30min of incubation on ice under agitation, the suspension (cells + Matrigel) was centrifuged at 1300rpm for 3min. RNA was extracted from the pellet using the Nucleospin kit (BioKé) protocol and sent to BGI for RNA-sequencing using the Illumina-HiSeq2500/4000 sequencer.

Raw fastq files received from BGI were processed using the following pipeline. Raw data quality control was performed using FastQC v.0.11.4^48^. Adapter removal and trimming of low-quality reads was done with Trimmomatic v.0.32^49^, and afterwards a custom Python script (ran with Python v.2.7.13) was used to remove reads with undetermined bases. Alignment of processed reads was performed using STAR v.2.5.3.a using GENCODE mouse release M15 (GRCm38)^50^. Aligned reads were then quantified using RSEM v.1.3.0^51^ and transcripts per million (TPM) were retrieved.

In order to identify genes showing interaction effect between cell lines and culture type, differential expression analysis was done using DESeq2 package for R software v.3.5.0^52^. Likelihood ratio tests were done for each gene fitting a model adjusted for cell line, culture type, batch, and the interaction effect.

A pre-ranked Gene Set Enrichment Analysis (GSEA, v2.2.07)^53^ was run with genes ranked using the negative logarithm (base 10) of the p-value and the sign of the beta coefficient from the differential expression model. For the interaction analysis, a positive interaction beta coefficient denotes a higher slope in D2OR. This can correspond to multiple biological behaviors: up-regulation in D2.OR 3D vs. 2D and down-regulation in D2A1 3D vs. 2D; up-regulation 3D vs. 2D in both D2OR and D2A1 but stronger in D2.OR; and down-regulation 3D vs. 2D in both D2.OR and D2A1 but stronger down-regulation in D2A1. Analogously, a negative interaction beta coefficient corresponds to the opposite behaviors. A total of 4018 gene sets were tested after filtering for minimum set size of 30 genes, including specific sets to assess dormancy^54^ and senescence^55^, but also more general gene sets, like hallmarks, canonical pathways (KEGG, Biocarta, Reactome, PID), and gene ontology, from MSigDB v.7.0^56^. For graphical representation, gene sets were clustered by gene overlap.

In order to assess intrinsic subtypes (PAM50) from the cell lines, we used Uniform Manifold Approximation and Projection (UMAP), a non-linear dimension reduction technique. To do so, we used primary breast cancer TCGA data (TPM downloaded from TCGA2BED)^57^ with available PAM50 information^58^ and computed a UMAP using a mouse-derived intrinsic subtype 1841-gene signature^31^, n = 50 neighbors and a minimal distance of 0.2.

### COBRA karyotyping

Metaphases from mouse cell lines were harvested using standard techniques, described in detail^59^. Multicolor fluorescence in situ hybridization (dubbed as COBRA-FISH) using mouse chromosome painting set to identify each chromosome was prepared and hybridized as presented in the detail published protocol^60^, following the procedure including image acquisition and software tools.

### Animal experiments

All animal experimental protocols were approved by the animal welfare committee of the Leiden University Medical Center and the Dutch Animal Experiments Committee. Female mice, aged between 8 and 12 weeks, were used exclusively.

Mice were injected with 1×10^6^ D2 H2B-Dendra2 cells loaded with DiI in the mesenteric vein. In short, mice received buprenorphine (0.01 mg/kg (Temgesic)) 30 minutes before surgery. Mice were anesthetized using injection anesthetics (70 mg/kg ketamine (anesketin) and 0.7 mg/kg medetomidine hydrochloride (sedastart) injected i.p.). Surgery was performed under aseptic conditions. A midline incision of ~1 cm was made in the belly region, the intestines were placed on a sterile gauze. Using a binocular, injection of tumor cells was performed in the mesenteric vein. Afterwards, the puncture wound was closed by pressure using a q-tip, and the animal closed using a suture. After surgery, mice received 1 mg/kg atipamezole hydrochloride (antisedan) s.c. to wake up. 2 weeks after injection, mice were sacrificed and the liver was harvested and fixed for 1 day in PLP fixation mix (1% paraformaldehyde, 0.2% NaIO_4_, 61□mM Na_2_HPO_4_, 75□mM L-Lysine and 14□mM NaH_2_PO_4_ in H_2_O). After fixation, tissues were incubated in 30% sucrose in PBS for > 6 hours, and frozen in tissue freezing medium (OCT). 10 μm sections were obtained at three different heights of the liver (at least 250 μm apart). Sections were embedded in vectashield (hardset with DAPI), and imaged on the slide scanner (3DHISTECH panoramic 250).

### Intravital microscopy

Mice were injected with 1×10^6^ D2 H2B-Dendra2 cells loaded with DiI in the mesenteric vein. 16 days later, the liver was surgically exposed and intravitally imaged on a Zeiss LSM 710 NLO upright multiphoton microscope equipped with a Mai Tai Deep See multiphoton laser (690–1040 nm) and a custom-fit stage with window holder. During imaging, the mice were anesthetized using isoflurane. Tumor cells visible underneath the window were recorded at multiple positions using z-stack timelapses. Images were collected in 8 bit and 512 x 512 pixels, with an xy-resolution of 0.25-083 μm and a z-resolution of 3 μm per pixel. Z stacks were recorded every 20 minutes for ~3 hours. Dendra2 and DiI were excited with 960 nm, and emission was collected in BiG NDD (510-550 nm), BiG NDD (575-620). Timeseries were registered using the ImageJ FIJI plugin Descriptor-based series registration (2D/3D + t). Maximum projections were made of 2 or 3 Z planes. Some images were smoothed and gamma corrected to enable visualization of lower intensity cells.

### Statistical Analyses

Data were analysed using software Graphpad Prism v5. Student’s t-test, Mann Whitney U test, or one- or two-way ANOVA statistical tests were used where appropriate. Statistical significance was defined as: n.s. P≥0.05, *P≤0.05, ** P≤0.001, *** P≤0.001, **** P≤0.0001. A *post hoc* analysis was only performed if a significant interaction was found. In the legend or figures, only significant analyses are shown, unless otherwise stated. Quantitative data are presented as the mean ± standard error of the mean, unless stated otherwise.

## ACKNOWLEDGEMENTS

We would like to acknowledge 1. all (former) members of the Ritsma and ten Dijke lab for help, support and critical comments; 2. the CCB microscopy facility for microscopy support; and 3. The LUMC FACS sort facility for flow cytometry support; 4. Willem Sloos for general lab/IT support. We also acknowledge our funders: CP was supported by Leiden University Profile area Bioscience: the science base of health ‘Building Pipelines for Drug Discovery’. LR was supported by a Gisela Thier fellowship from the Leiden University Medical Center (LUMC), a Veni grant from the Netherlands Organisation for Scientific Research (NWO, 016.176.081) and a subsidy from the Leids Universiteits Fonds (LUF, CWB 7204). CP, LR and PtD members were supported by a subsidy from the Cancer Genomics Center Netherlands. XS was supported by by the Spanish Ministry of Science and Innovation, RTI2018-102134-A-I00 (MCIU/AEI/FEDER, UE).

## AUTHOR CONTRIBUTIONS

LR conceived the study. LR and CP designed the study. LR, CP, KA, DLM, MvD, AT, SvG, AT, AGJ, KS performed experiments. AA, RS performed bioinformatic analyses. CP and LR wrote the manuscript. LR, PtD, XS supervised the study. All authors approved the final version of the manuscript.

## COMPETING INTERESTS STATEMENT

The authors declare no competing interests.

## SUPPLEMENTARY FIGURES AND LEGENDS

**Supplementary Fig. 1:**
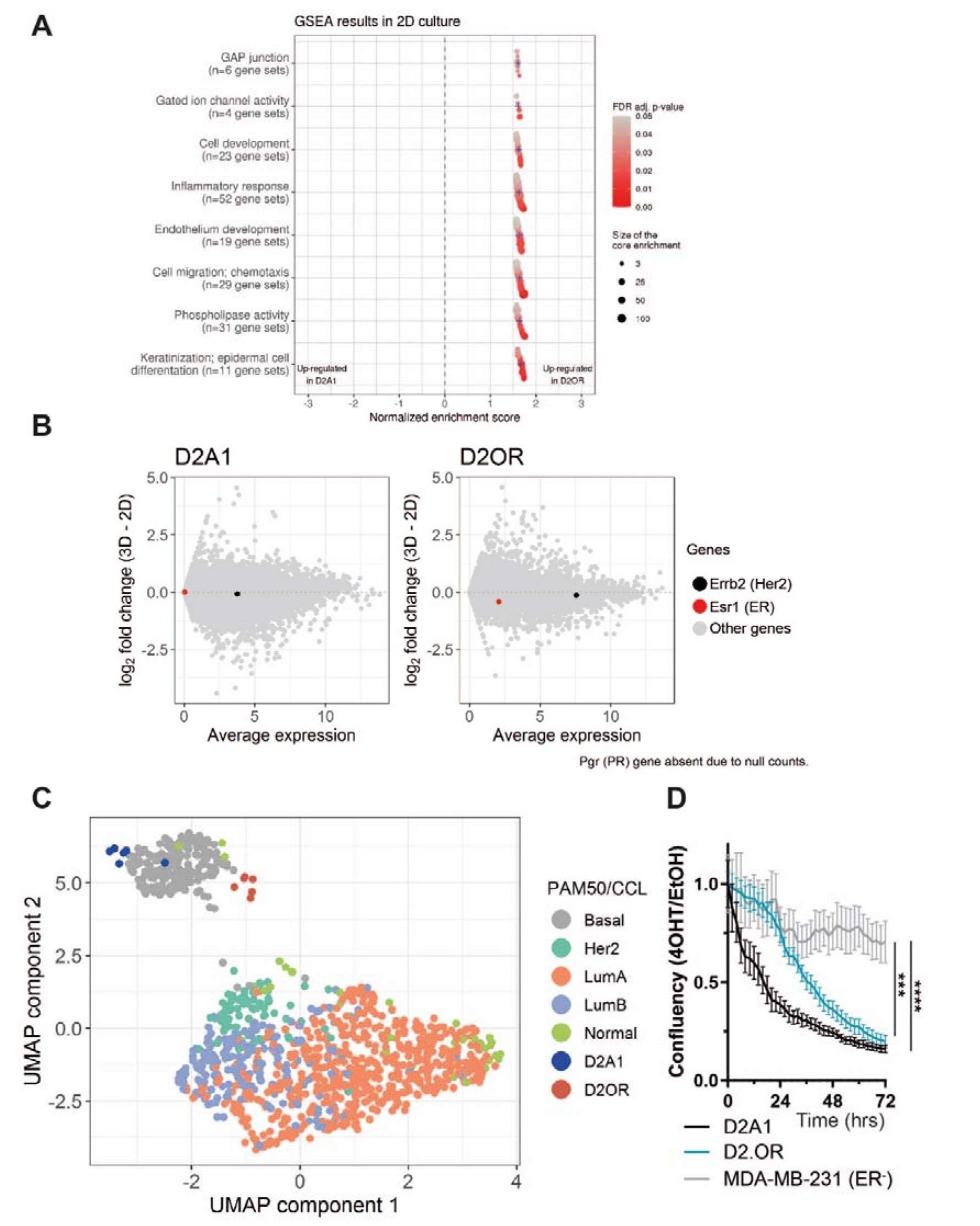
D2 cells are considered to be of the ER+/HER2+ basal subtype. **a** GSEA analysis on the differential expression between D2A1 and D2.OR cultured in 2D. Gene-sets ordered by normalized enrichment score (NES) within each cluster. Blue crosses represent average NES in each cluster. Point color represents FDR-adjusted p-value and size represents core enrichment. **b** MA plots for D2A1 and D2.OR cells (i.e., x-axis: average expression in 2D and 3D; y-axis: log_2_ fold change between 3D and 2D). Erbb2 and Esr1 are highlighted. **c** Combined UMAP analysis of D2A1 and D2.OR transcriptional data (in 2D) and TCGA primary breast cancer dataset with available PAM50 information, using a mouse-derived intrinsic subtype 1841-gene signature. **d** Quantification of cell viability in 2D treated with 4OHT [5μM] or vehicle as measured by PrestoBlue assay. Data is normalized to day 1, the ratio of 4OHT over vehicle is plotted. Time x Cell line, F (72, 216) = 16.36, P = ****. *Post hoc* test, MDA-MB-231 vs D2A1 or D2.OR P = ****. P value was calculated using 2-way RM ANOVA.

**Supplementary Fig. 2:**
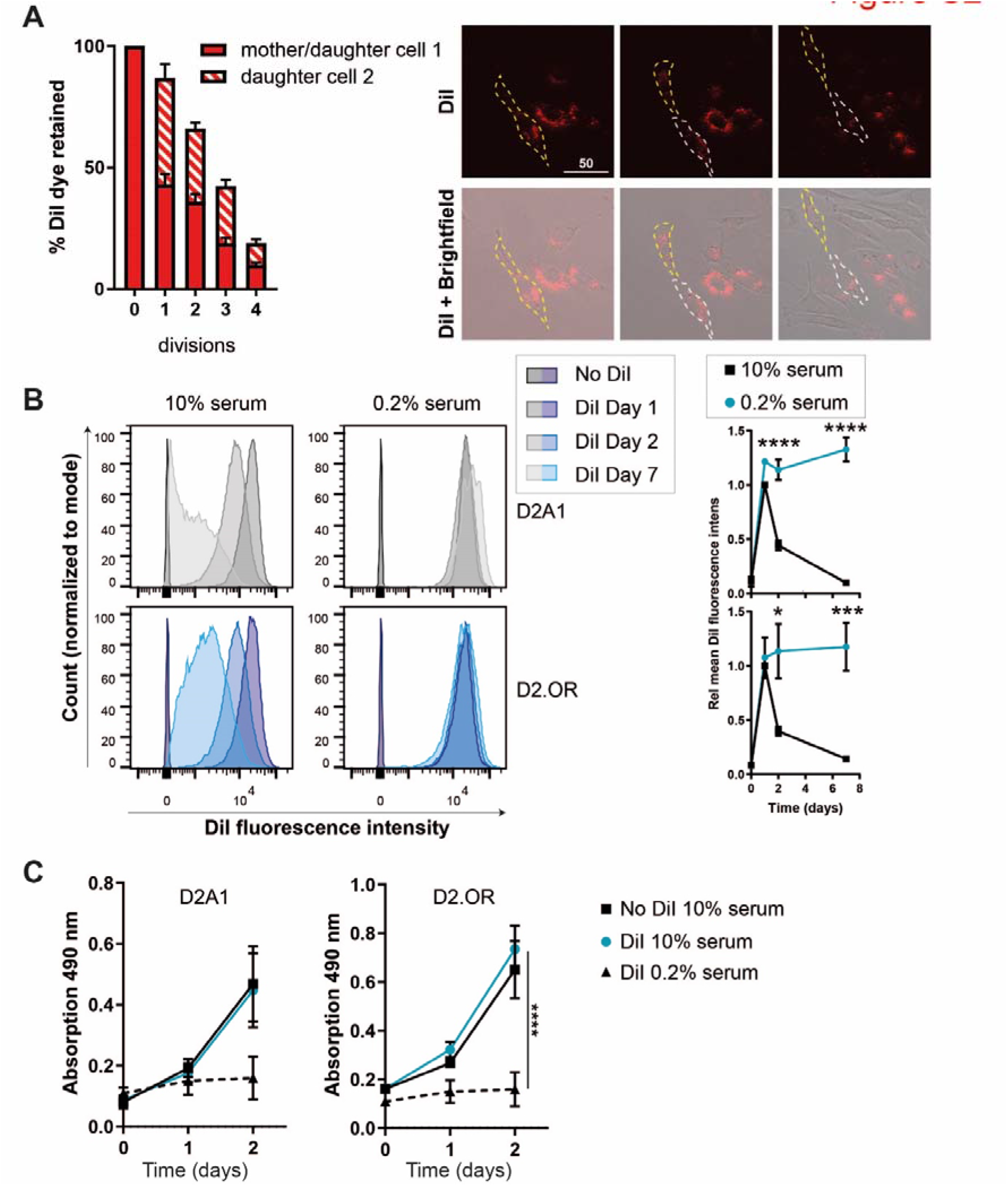
The effect of DiI loading of D2 cells. **a** Quantification (left) of live cell imaging time series (right) of D2A1 cells in 2D. Individual cells were tracked and total DiI fluorescence intensity per cell was determined (background subtracted) for mother cells (yellow dashed line) and subsequent daughter cells (white / yellow dashed line). N = 5 cells, scalebar, 50 μm. **b** FACS plots depicting D2 cells loaded with DiI and cultured (in 10 or 0.2% FBS) for 1, 2 or 7 days. Quantification on the right. N = 3. Interaction D2A1 (time x serum), F (3, 14) = 5.470. P = *. Interaction D2.OR (time x serum), F (3, 14) = 38.12, P = ****. **c** Quantification of MTS assay of D2 cells loaded with or without DiI, cultured in 10% or 0.2% FBS for 0,1, and 2 days. N = 3. Interaction D2A1 (time x condition), F (4,12) = 3.426, P = *. Interaction D2.OR (time x condition) F (4,12) = 5.144, P = **. P value was calculated using 2-way RM ANOVA.

**Supplementary Fig. 3:**
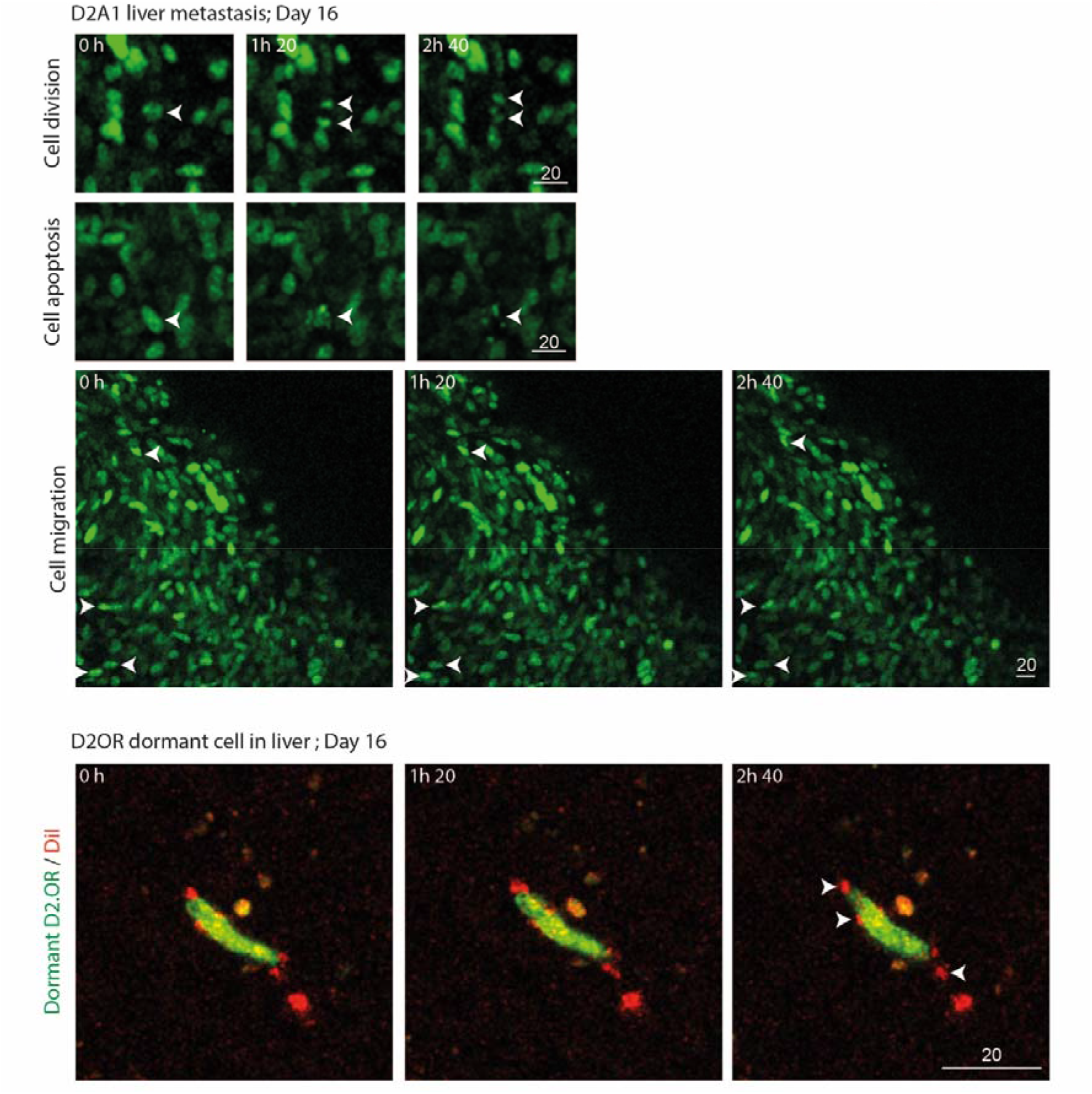
Intravital imaging of D2A1 and D2.OR cells in the liver. D2A1 and D2.OR H2B-Dendra2 cells were intravitally imaged 16 days after injection in the mesenteric vein for ~3 hours. In D2A1 liver metastases cell division, apoptosis and cell migration was observed. D2.OR cells were loaded with DiI before injection, which was retained at day 16. During 3 hour movies, the cells showed no signs of division, apoptosis or migration. Arrowheads indicate D2A1 cell division/apoptosis/migration or DiI dye in a D2.OR cell.

**Supplementary Fig. 4:**
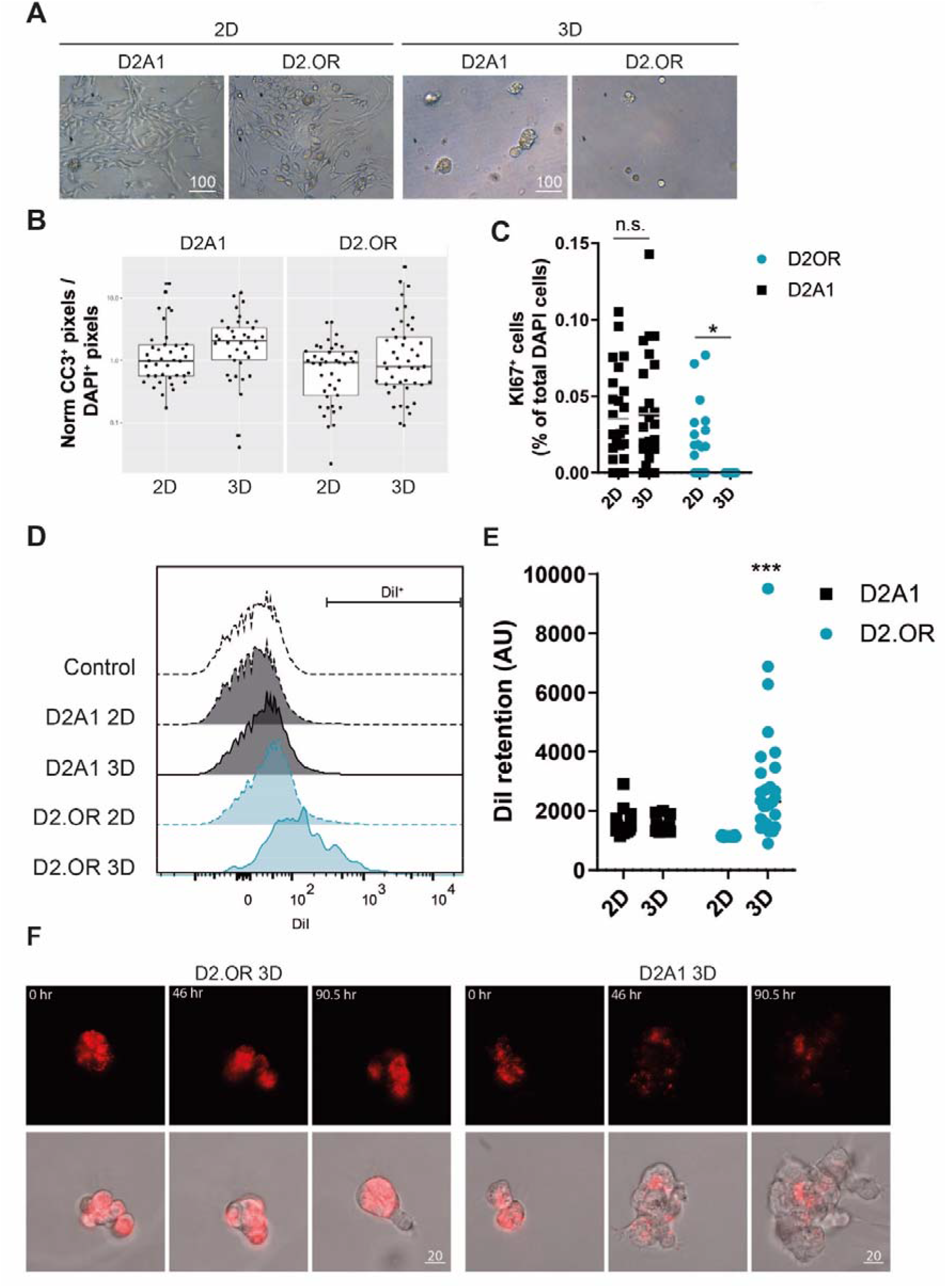
D2.OR cells are dormant in an *in vitro* 3D condition. **a** Images of D2 cells cultured in 2D or 3D for 6 days. Scalebar, μm. **b** Quantification of immunostaining of D2 cells cultured for x days; per FOV cleaved caspase 3 (CC3)^+^ pixels were divided to DAPI^+^ pixels to account for the number of cells, and normalized per replicate to D2OR 2D. N ≥ 37 FOV in 4 replicates. ANOVA not significant. **c** Quantification of immunostaining of D2 cells cultured for x days; percentage of KI67+ cells per FOV. Interaction, *F* (1,136) = 4.696, P = *. *Post hoc* test results are shown in figure. N = 25 FOV of 4 replicates. **d** FACS plot related to figure **2b**. **e** Quantification of DiI fluorescence in microscopy tile scans of D2 cells loaded with DiI grown in 2D or 3D for 7 days. N ≥ 20 FOV. Interaction, F (1,85) = 13.49, P = ***. *Post hoc* test, D2.OR 3D vs D2.OR 2D, P = ****. D2.OR 3D vs D2A1 2D or 3D, P = ***. **f** Microscopy images from a time series of D2 cells loaded with DiI and grown in 3D for 4 days, imaged every 30 mins. Upper panels; DiI. Lower panels; DiI and brightfield. Scalebar, μm. P values were calculated by 2-way ANOVA.

**Supplementary Fig. 5:**
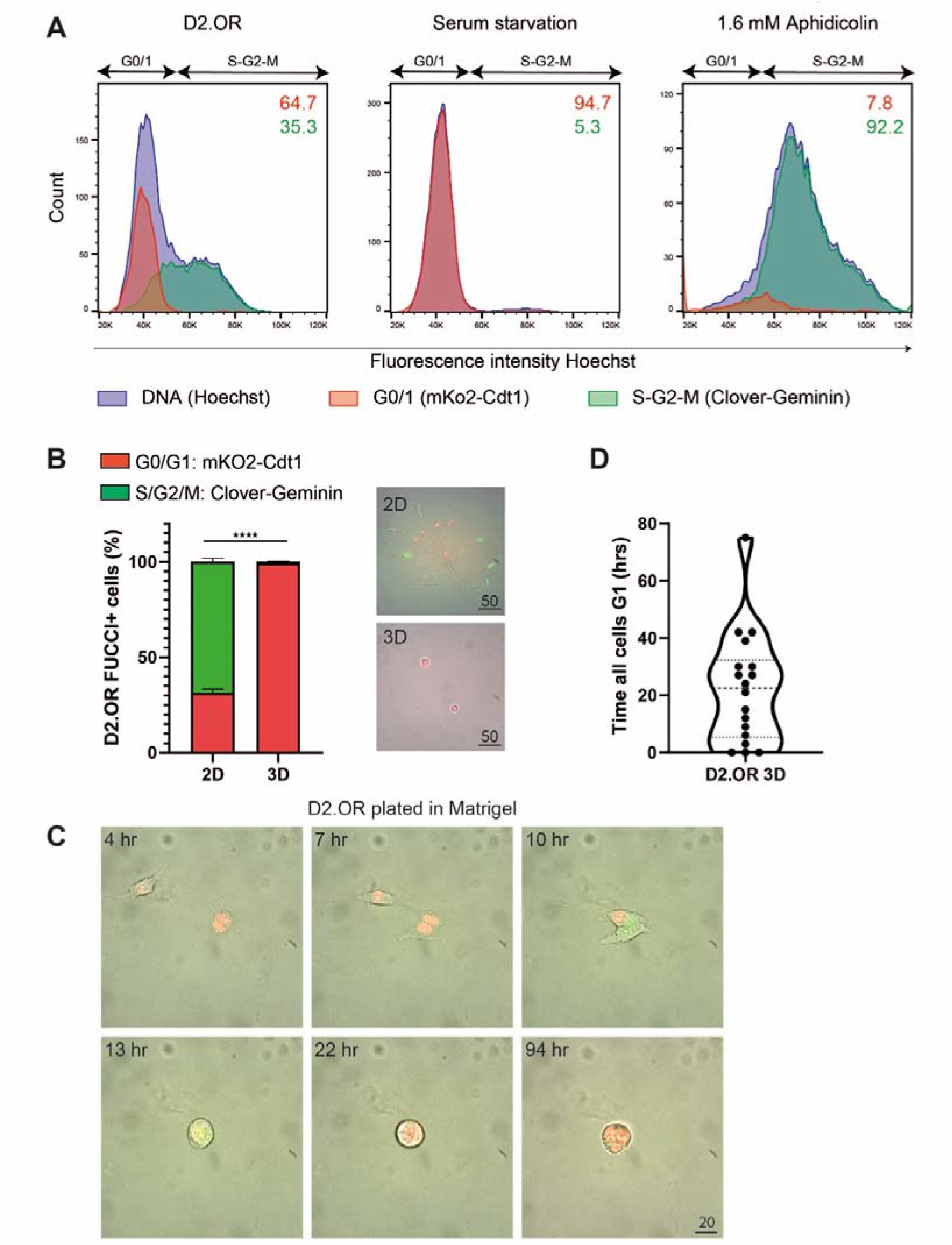
Characterization of D2.OR Fucci cells. **a** FACS plots depicting cell cycle status (based on Hoechst) of D2.OR cells in 2D treated with serum starvation (0.1% FBS) or aphidicolin [1.6mM] for 24 hrs. The percentage of mKo2 (red) or Clover (green) cells based on the Hoechst population is shown in the top right corner. **b** Quantification (left) of Fucci mko2^+^ or Clover^+^ cells in microscopy images (right). D2.OR Fucci cells were cultured for 2 days in 2D and for 7 days in 3D. Interaction effect, F(1,20) = 679.7, P = ****. *Post hoc* test, Clover vs mKO2, P = ****. **c** related to **Movie 2**. Images from time recordings of D2.OR Fucci cells plated in Matrigel. The time indicates the time after plating in Matrigel. Images were taken every 3 hours. **d** Related to **fig. 2d**. D2.OR Fucci cell clusters in 3D were analyzed in time lapse movies to determine the time until all cells in that cluster were mKo2^+^. N = 18 clusters. P value was calculated by two way ANOVA.

**Supplementary Fig. 6:**
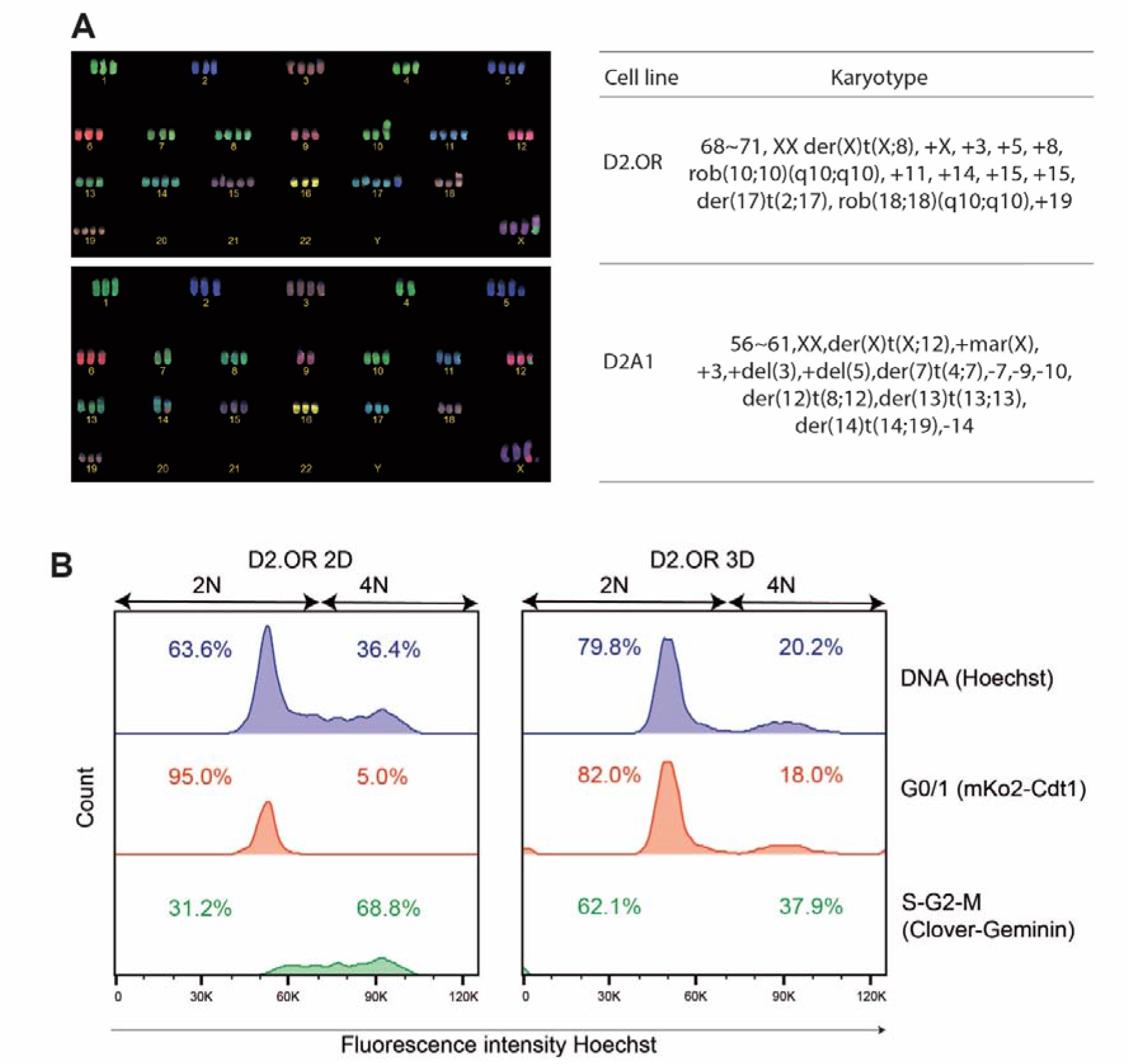
D2 cells are polyploid. **a** Cobra karyotyping of D2.OR and D2A1 cells in 2D. **b** FACS analysis related to **Fig. 2f**. Cell cycle analysis based on Hoechst is shown. Percentages indicate cells in 2N or 4N gate for that color.

**Supplementary Fig. 7:**
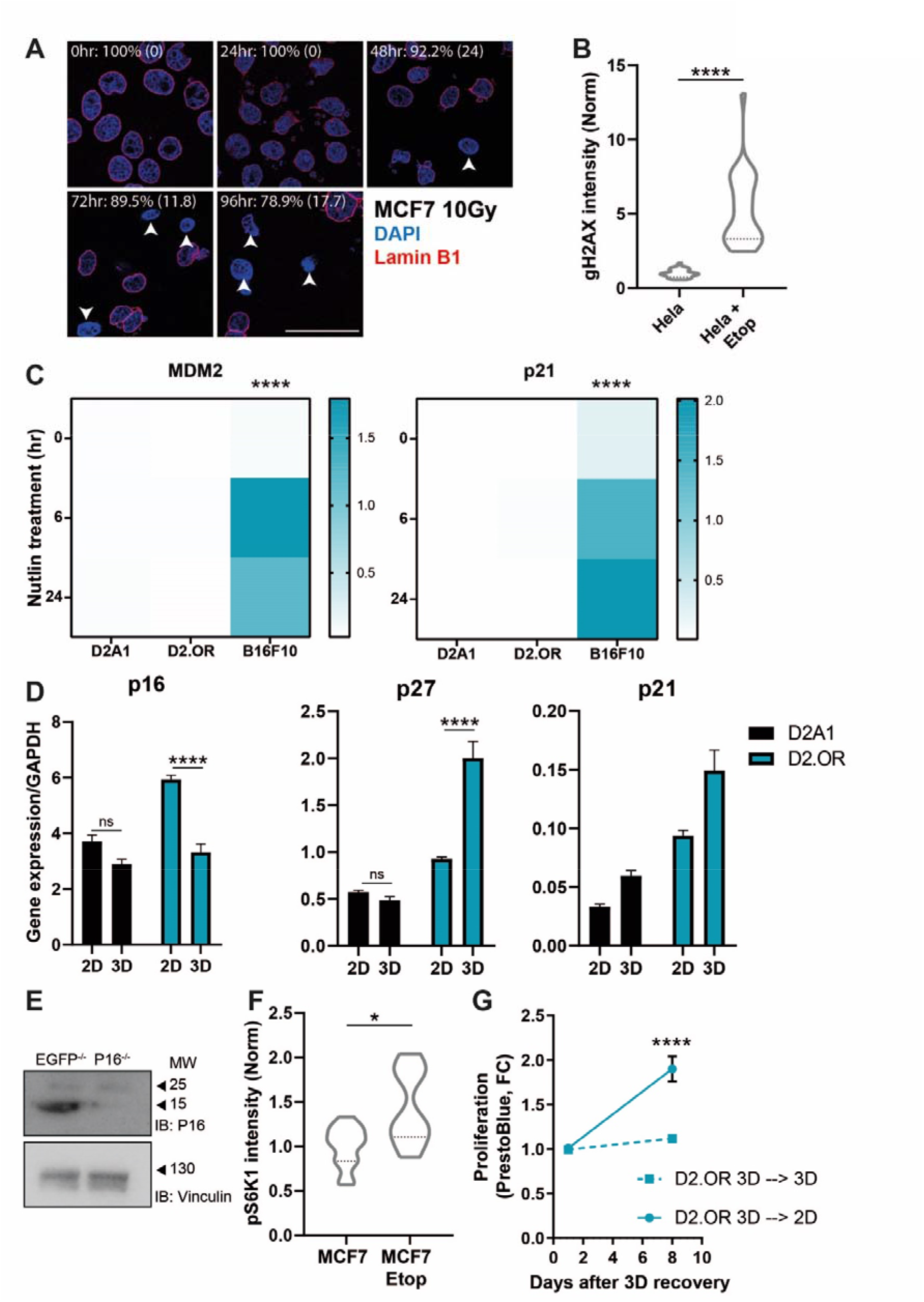
Characterization of dormancy and senescence parameters. **a** Microscopy images of MCF7 cells cultured in 2D, irradiated with 10 Gy (1Gy/min), and incubated for indicated time to induce senescence. Cells were then stained for DAPI and LaminB1. Percentage of positive laminB1 cells (±SD) is indicated. N = 10 FOV per condition. Scale bar, 50 μm. **b** Related to **Fig. 4c**. Quantification of nuclear γH2AX staining intensity of Hela cells cultured for 4 days in 2D. 24 hrs before fixation cells were treated with Etoposide (12.5 μM) to induce senescence. Replicates normalized to Hela. N = 32 field of view (FOV) of 3 replicates. **c** Heatmap showing qPCR normalized gene expression of D2A1, D2.OR and B16F10 (+ control) cells in 2D treated with Nutlin-3 [10 μM] for 0, 6 or 24 hours. Interaction MDM2, F (4,18) = 426.9, P = ****. *Post hoc* test, B16F10 vs D2.OR or D2A1, P = ****. Interaction p21, F (4,18) = 46.75, P = ****. *Post hoc* test, B16F10 vs D2.OR or D2A1, P = ****. **d** Plots showing normalized gene expression of D2 cells in 2D or 3D cultured for 7 days. Interaction p16, F(1,11) = 14.80, P = n**. *Post hoc* test, D2.OR 2D vs 3D, P = ****. Interaction p27, F(1,12) = 41.83, P = ****. *Post hoc* test, D2.OR 2D vs 3D, P = ****. Interaction p21, F (1,12) = 2.455, P = n.s. N = 2 replicates in duplo. **e** Immunoblot showing lysates of D2.OR EGFP-/- and D2.OR p16-/- cells cultured in 2D. **f** Quantification of pS6K1 immunostaining of MCF7 cells cultured in 2D and treated with or without etoposide (12.5 μM) (senescence control). Replicates are normalized to MCF7 vehicle. N ≥ 10 field of view (FOV) of 2 replicates. **g** Quantification of D2.OR cell proliferation after extracting the cells from 3D and re-plating them into 2D or 3D. Time x cell, F (1,6) = 30.24, P = **. *Post hoc* test, 3D vs 2D day 8, P = ****. N = 2 in duplo. P value was calculated by Mann-Whitney or two way ANOVA test.

**Movie 1: Live cell imaging of D2.OR Fucci cells plated in Matrigel.** D2.OR Fucci cells were plated in Matrigel. Time indicates time after plating. Images are a merge of brightfield, Fucci mKO2 (Red) and Clover (Green) taken every 3 hours.

**Movie 2: Live cell imaging of dormant D2.OR Fucci cells extracted from Matrigel and replated in 2D.** D2.OR Fucci cells were cultured in Matrigel for 4 days. Then, the cells were extracted from the Matrigel and replated on plastic μ-ibidi dishes and imaged every 20 minutes for 3 days.

**Supplementary table 1.** Gene sets included in GSEA analysis comparing D2OR and D2A1 in 2D.

**Supplementary table 2.** Gene sets included in GSEA analysis comparing the interaction effect when comparing D2A1 and D2.OR in 2D and 3D.

## Notes

### Competing Interest Statement

The authors have declared no competing interest.

